# Drivers of tandem repeats variation within and between stick insect genomes

**DOI:** 10.64898/2026.09.22.753450

**Authors:** William Toubiana, Marion Leleu, Vincent Mérel, João Souto, Marie Raynaud, Tanja Schwander

## Abstract

Tandemly repeated sequences are noncoding, highly variable, yet integral components of eukaryotic genomes. Empirical research has so far largely focused on a few well-characterized, functional tandem repeats (TRs) or documented the broad diversity and rapid evolution of these repeats without explicitly testing the forces driving their evolution. By characterizing TR abundance and TR sequence composition within and between chromosomes and species in the stick insect genus *Timema*, we uncover that TR loads are enriched in genomic regions of high recombination but tend to be reduced in species with higher effective population sizes. These results suggest that there is a mutational effect of recombination that generates increased TR loads and that the bulk of TRs experience weak selection, leading to their rapid turnover over short evolutionary timescales. Finally, comparisons of centromere and non-centromere TRs reveal that the rapid divergence of centromeres is best explained by the combination of the fast-evolving nature of TRs *per se* and centromere-specific evolutionary forces. Overall, this study provides the first in-depth investigation of genome-wide TR evolution, while integrating the effects of recombination, genetic drift, and centromeric position along the genome.

## Introduction

Tandem repeats (TRs), also referred to as satellite repeats, are DNA sequences repeated in a tandem fashion (i.e., head-to-tail arrangement) and that form arrays that may span several megabases ((1); Figure 1A)). They are an integral component of eukaryotic genomes, notable for their high variability in copy number and sequence composition (2). However, the evolutionary forces driving their maintenance and diversity within and between species are still unclear and debated. Central to this debate is the extent to which TRs serve biological functions, and whether they should be considered largely non-functional or mostly functional elements of the genome (3, 4). Specifically, well-known TRs such as ribosomal DNA (rDNA) or telomere repeats are essential for processes related to protein biosynthesis and chromosomal integrity (5, 6). However, whether such functional TRs are rare exceptions or representative of the TRs overall remains unknown. Indeed, the inherent propensity of TRs to form and expand, along with their substantial variability across taxa, argues for the bulk of them being non-functional (3).

**Figure 1:**
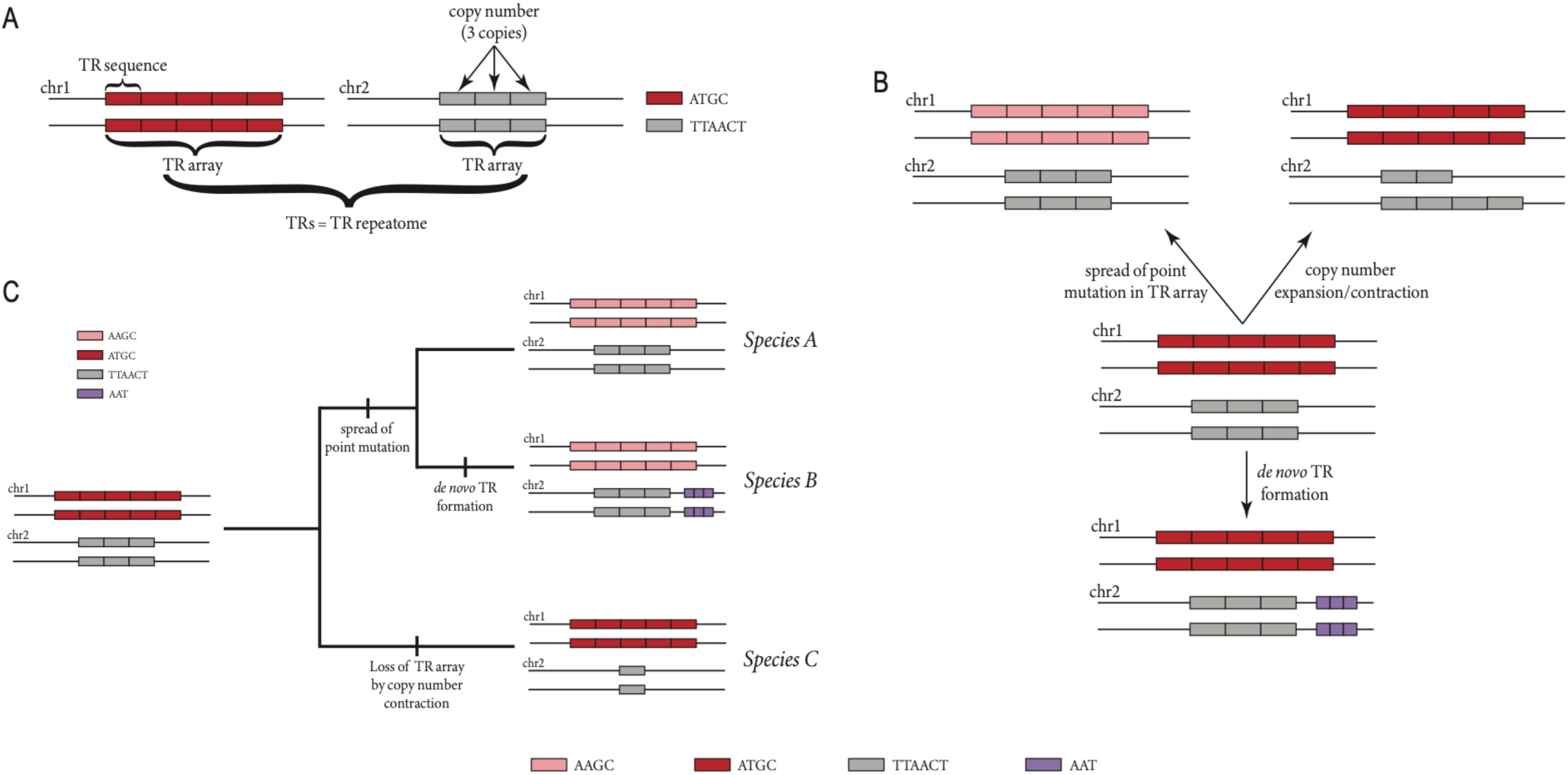
A) Terminology used in the present study and structure of tandem repeats (TRs). The TR sequence represents a DNA motif which is repeated in tandem fashion (head to tail arrangement). The set of consecutive repeats forms a TR array, which can vary in copy number and DNA composition of the TR sequence (as shown in the schematic). The total collection of all TRs across the genome then constitutes the TR repeatome. For simplicity, the genome-wide TRs are here represented on two chromosomes (chr1 and chr2) and by one TR array per chromosome. B) Mutational processes driving TR repeatome variation over time. First, a point mutation can occur within a TR array (e.g., change of “AAGC” to “ATGC” TR sequence) and subsequently spread to the extent that the formal TR sequence is no longer observed in the array. Second, the addition or removal of copies from an existing TR array results in its expansion or contraction, respectively. Third, a new TR array can emerge from a single-copy sequence (e.g., “AAT” sequence) and lead to de novo TR formation. C) TR repeatome turnover among species. As a consequence of the three mutational processes described in B), species can share or lack specific TR sequences. Some initially shared TR sequence may become species-specific, as with the “AAT” and “ATGC” TR sequences in Species B and C, respectively. Note that in Species C, the “TTAACT” sequence is no longer considered a TR sequence due to its copy number contraction that resulted in the loss of the TR array.

Theoretical studies have predicted TR evolution under a non-functional framework, examining how the loads of a specific repeat vary with recombination, genetic drift, and selection (7). Within this framework, TRs are considered slightly deleterious elements for the host, becoming harmful as they accumulate to excessively high levels (8). Consequently, TRs are predicted to accumulate in species with small effective population sizes (*Ne*), where the effect of drift is stronger and the efficacy of selection against excessively high TR loads is reduced (8). Increased recombination is also expected to impact TR load by generating a wider distribution of copy numbers in the population through unequal crossovers (8). This process, along with recombination’s classical role in reducing linkage between genetic loci (which increases *Ne* (9, 10)), further enhances the efficacy of selection against high TR loads. While these theoretical models offer testable predictions on the evolution of genomic TR loads, empirical studies typically focus on specific (often functional) TRs or describe the overall set of TRs without explicitly testing the evolutionary forces underlying their load variation within or between genomes ((11-18) but see (19, 20)).

Beyond changes in global and local genomic loads, TRs also evolve at the sequence level, resulting in variable genome-wide repertoires of TRs in different species (i.e., TR repeatome, also referred to as satellitome (21); Figures 1B and 1C). TR repeatome evolution is driven by three major processes 1) point mutations within individual TR sequences, which sometimes spread throughout TR arrays and replace ancestral TR variants, 2) *de novo* emergence of new TRs, 3) copy number expansions and contractions that lead to the gain or loss of TR arrays ((22, 23); Figures 1B and 1C). These mutational processes are then modulated by recombination, drift and selection, which collectively contribute to the overall difference (“turnover”) in TR repeatomes between species. Although no existing evolutionary model predicts how the TR repeatome should evolve, considering the joint mutational processes and different selective regimes, its rate of turnover (i.e., the proportion of non-identical TR sequences as a function of species divergence) is expected to reflect the functionality of TR sequences. Specifically, if the majority of TR sequences are functional, the TR repeatome turnover should be slow due to purifying selection constraining the evolution of individual TR sequences (24, 25). By contrast, if most TR sequences are non-functional, the TR repeatome turnover between species could be fast, driven by recurrent loss or *de novo* formation of TRs, and spread of point mutations across arrays (see Figure 1). Paradoxically, certain TRs such as those composing centromeric regions (i.e., centromeric TRs) are well-known to evolve rapidly despite their essential role in chromosome segregation (26). These rapid changes are often attributed to centromere drive during female meiosis (26). Yet, whether centromeric TRs exhibit distinct, and markedly faster turnover dynamics relative to other TR sequences in the genome remains as an open question.

In this study, we investigate the load and sequence variation of the TR repeatome in relation to varying levels of recombination, genetic drift, and selection within the stick insect genus *Timema*. *Timema* offers a compelling system for this purpose owing to its extensive variation in effective population size among closely related sexual species, in addition to recombination rate variation within species/genomes, due to chromosome number (XX:X0 sex determination system) and chromosome size (27, 28). Through a first comparative approach on TR load across species, we show that the proportion of TRs in *Timema* genomes tends to be negatively correlated with effective population size. This aligns with theoretical models that consider TRs as slightly deleterious elements. Within genomes however, we find an enrichment of TRs in regions of high recombination, likely due to recombination generating *de novo* TRs as well as extended TR arrays. In a second approach, we investigate how the level of TR repeatome turnover scales with phylogenetic distance between species and differs from genomic regions subject to variable selective constraints, such as coding DNA sequences (CDS), introns, and centromeres. Our results support the view that the vast majority of TR sequences experience weak selection, leading to their rapid turnover over short evolutionary timescales. Moreover, we find that this intrinsically fast-evolving nature of TRs must be considered alongside centromere-specific features, such as centromere drive, when assessing the evolutionary forces shaping centromere evolution.

## Results

### Variation in TR loads, *Ne* and genome sizes across sexual *Timema* species

To estimate TR load variation within and between five sexually reproducing *Timema* species, we annotated TRs in five available nanopore-based genome assemblies. We then mapped whole-genome Illumina reads of 4 to 7 females per species onto these TR-annotated genome assemblies (see details in Material and methods, Supplementary table 1). For the re-sequenced samples, we estimated genome-wide TR load as the proportion of the genome covered by annotated TRs. This proportion was calculated as the ratio of mean per-base read depth across TR-annotated regions to mean genome-wide read depth, thereby accounting for individual differences in sequencing depth. The resulting estimates were highly consistent with those obtained from TR array lengths in the reference assemblies (see details in Material and methods). The genome-wide proportion of TRs varies considerably between species, ranging from 8% to 17% of the total DNA content, with relatively little variation within species (Figure 2A). We also find that variation in TR load explains most of the genome size variation in *Timema*, with 90-95% of genome size variation across species explained by variation in TR proportion (Figure 2A; adjusted R^2^ = 0.95, p-value = 0.003; Phylogenetic Generalised Least Square: adjusted R^2^ = 0.90, p-value = 0.009).

**Figure 2:**
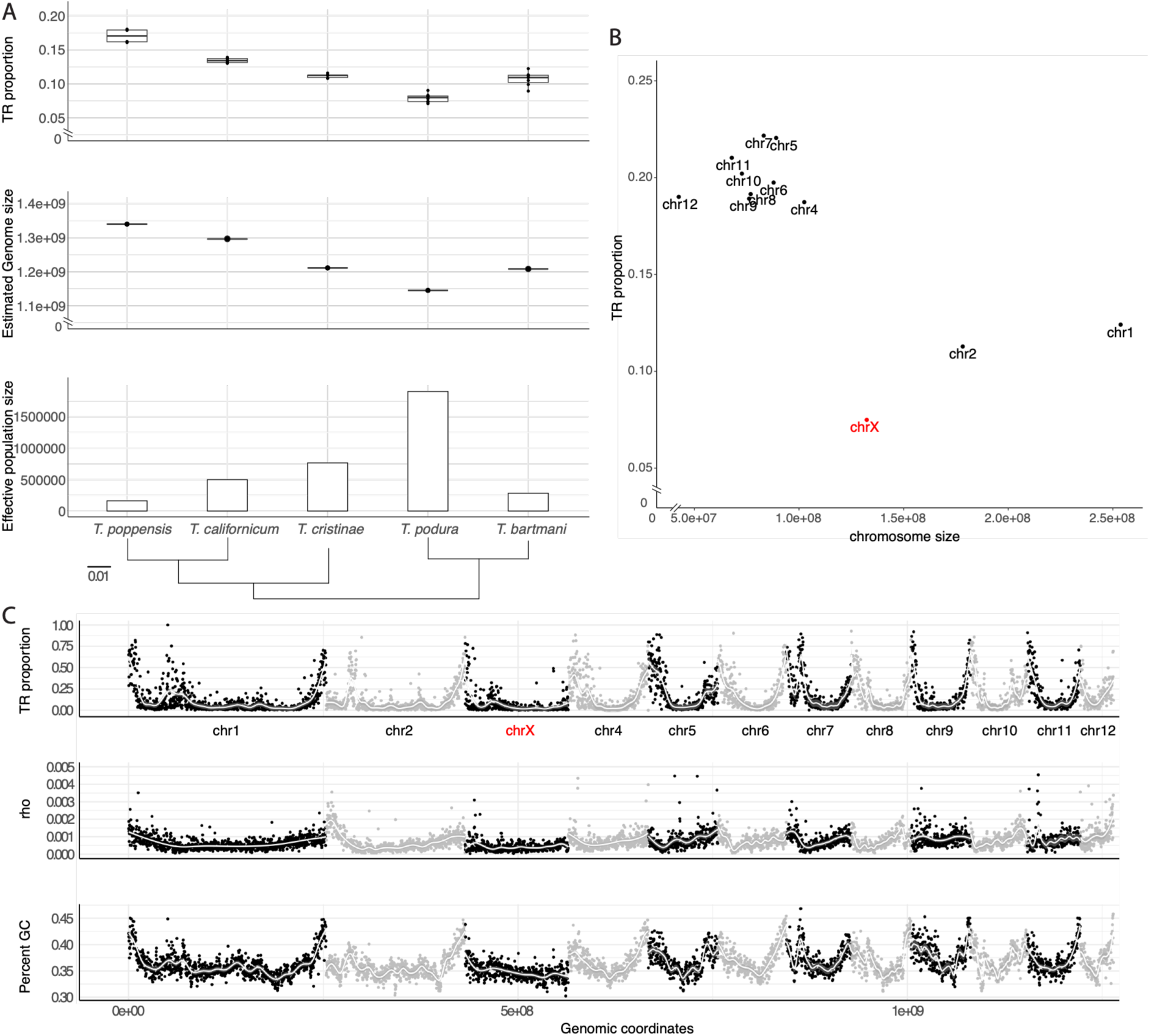
Tandem repeat (TR) load variation within and between genomes. (A) Comparison of genome-wide TR load with genome size and effective population size across five sexual Timema species. The phylogenetic tree was inferred from whole-genome sequence data (27) and branch lengths represent genetic distances (i.e., substitutions per site). Each boxplot is based on 4 to 7 females per species. (B) Relationship between chromosome size and TR proportion in T. poppense. (C) Chromosomal distribution of (from top to bottom) TR proportion, recombination rate (rho values), and GC content across the 12 chromosomes of T. poppense. For visualization purposes, the highest rho values on chromosomes 4, 9 and 11 are not shown. Alternating black and grey shading distinguishes individual chromosomes. Each point represents a non-overlapping 250-kb genomic window.

We previously estimated, based on nucleotide polymorphism data, that the five *Timema* species under study are characterized by up to 10-fold variation in effective population size (27). This variation is expected to influence genome evolution, including the accumulation of TRs, by modulating the action of selection relative to drift (i.e., the efficacy of selection), although the power to detect such an influence is very low with n=5 species. Consistently, we identified a negative but non-significant relationship between TR proportion and effective population size estimates (Figure 2A; adjusted R^2^ = 0.48, p-value = 0.12 / Phylogenetic Generalised Least Square: adjusted R^2^ = 0.38, p-value = 0.16). Taken together, our data suggest that TRs accumulate and contribute to genome size expansion in *Timema*, perhaps particularly in species with smaller effective population sizes where the selection against slightly deleterious TRs is less effective.

### TRs accumulate in high-recombination regions within genomes

Within genomes, the efficacy of selection is expected to vary among chromosomes through variation in census sizes and recombination rates. Specifically, under XX/XO sex chromosome systems, the census size of the X chromosome population is 3/4 the size of an autosomal one. Furthermore, recombination is reduced on the X relative to autosomes (because in *Timema* the X only recombines in females, while autosomes recombine in both sexes (27)), as well as on large relative to small autosomes (because the typical single chiasmata per chromosome or chromosome arm leads to higher chromosome-scale recombination rates for small chromosomes (29, 30)). We examined whether this variation in selection efficacy co-varies with TR loads between chromosomes. Specifically, with a negative effect of selection on TR loads, we expected a positive relationship between autosome length and TR proportion and a higher TR proportion on the X chromosome compared to the autosomes. However, we observed the reverse pattern, with lower TR proportions on larger autosomes and the X chromosome across the examined *Timema* species (Figure 2B; Supplementary figure 1; adjusted R^2^_autosomes_ = 0.67, p = 0.001). The X chromosome notably comprises two times less TRs than expected given its size: all else equal, the X chromosome of *T. poppense* was expected to comprise about 15% of TRs while only 7.5% were estimated (Figure 2B). Together, our findings suggest that TR load variation among chromosomes is not strongly influenced by the effect of selection. Instead, we hypothesize that the pattern observed may be induced by a biased mutational effect of recombination which would generally increase TR loads.

To further investigate this, we examined fine-scale recombination rate variation within chromosomes (using 250-kb non-overlapping windows), using recombination landscapes inferred from population-level linkage-disequilibria in *T. poppense* (see Material and methods). Consistent with recombination contributing positively to TR loads within genomes, TR proportions are high in genomic regions (i.e., windows) where the recombination rate is elevated, most notably on chromosome ends (Figure 2C). Thus, we found a positive correlation between TR proportion and recombination rate (Spearman’s rho = 0.23, p-value < 0.001). Moreover, about 8% of the variation in TR proportion is explained by fine-scale recombination rate variation, once size differences between chromosomes were accounted for (Figure 2C; R^2^ = 0.08, p-value < 0.001).

We corroborated the inferences based on recombination landscapes by repeating the same analyses using GC content within windows as a proxy for recombination rates (31). As anticipated, we first observed that *Timema* chromosomes comprise regions (notably chromosomal ends) with elevated GC content, similar to variation described in other organisms (32), and which coincide with regions of high recombination (Figure 2C). These GC-rich DNA windows also correspond to regions of high TR loads, as evidenced by a strong and positive correlation between TR proportions and GC content (Figure 2C; Spearman’s rho = 0.75, p-value < 0.001). Furthermore, 55% of the variation in TR proportion can be explained by GC content (R^2^ = 0.55, p-value < 0.001).

Together, our results in *Timema* highlight a contrasting pattern on the TR load distribution between species and within genomes. While the TR load most likely increases as the effective population size becomes smaller and the selection efficacy reduces at the species level, TR loads are reduced in chromosomes and genomic regions of lower recombination rate. We propose that these opposing patterns are best explained by genome-wide purifying selection acting against TR accumulation, while the mutagenic effects of recombination locally promote TR expansions. The balance between these forces ultimately shapes the distribution and abundance of TRs within and among species.

### TR sequence length and array density are elevated in regions of high recombination

The specific mechanism through which recombination would cause a local TR load increase is unknown, although it likely results from the independent or combined increase in TR copy number, sequence length and array density (i.e., the number of TR arrays per chromosome or window; see Figure 1). To evaluate how recombination influences these three components, we analyzed how they vary with our two proxies of recombination rate, rho values and GC content. We find that differences in recombination rates within and between chromosomes have a strong positive influence on all three components (Supplementary figure 2). Specifically, a positive and significant association is detected between both proxies of recombination and TR copy numbers, sequence lengths, or array densities. Collectively, these results suggest that the proportion of TRs increases in regions of high recombination via multiple effects. First, recombination promotes the formation of TR arrays, consistent with previous theoretical simulations (23, 33). Second, our results indicate that recombination increases the proportion of TRs by generating longer TR sequences. Although the underlying mechanisms remain unclear, this may involve processes such as higher-order repeat formation (33). Finally, recombination also produces a net increase in TR copies within an array.

### TR repeatome turns over rapidly and is composed of chromosome-specific sequences

TR repeatomes vary substantially among species, not only in their overall abundance but also in their sequence composition. Differences in repeatome composition are commonly described as repeatome turnover because they do not correspond to a well-defined evolutionary process. Indeed, current analyses can only quantify the extent to which particular TR sequences are shared between species because they cannot distinguish the relative contributions of sequence divergence within ancestral repeats from copy-number changes leading to the complete loss or emergence of novel repeats. This limitation arises because TR repeatomes vary considerably over short evolutionary timescales and because no generally accepted model of TR sequence evolution exists. As a result, homology relationships among TRs become difficult or impossible to establish.

While considering the limitations imposed by the biology and theoretical knowledge of TR repeatome evolution, we evaluated the selective constraints that may be operating on TR sequences genome-wide by estimating their rate of turnover among *Timema* species and comparing it to the turnover of exon and intron sequences. Purifying selection is expected to constrain the turnover of TR sequences serving specific functions (18, 25, 34). Specifically, we quantified the proportion of shared sequences between *T. poppense* and the four other studied *Timema* species (covering 7 to 35 million years of divergence) as well as with its sister parthenogenetic species, *T. douglasi*, which diverged recently (35, 36). As expected, TRs exhibit a higher rate of turnover than exon sequences, with about 40% of TR sequences shared after a few hundred thousand years and only 15% retained after about 7 million years (Figure 3A). In comparison, homologous exon sequences that size-match the distribution of TR sequence lengths in the *T. poppense* genome (see details in material and methods), still show 75% of sequence sharing after a few hundred thousand years or nearly 50% after 7 million years (Figure 3A). Homologous introns show a turnover rate approaching that of TRs, with only about 20% retention after 7 million years (Figure 3A) but a somewhat stronger short-term conservation (with 60% sharing after a few hundred thousand generations of divergence). Overall, these patterns of turnover support the general idea that the bulk of TR sequences varies rapidly and evolves under weak selective constraint.

**Figure 3:**
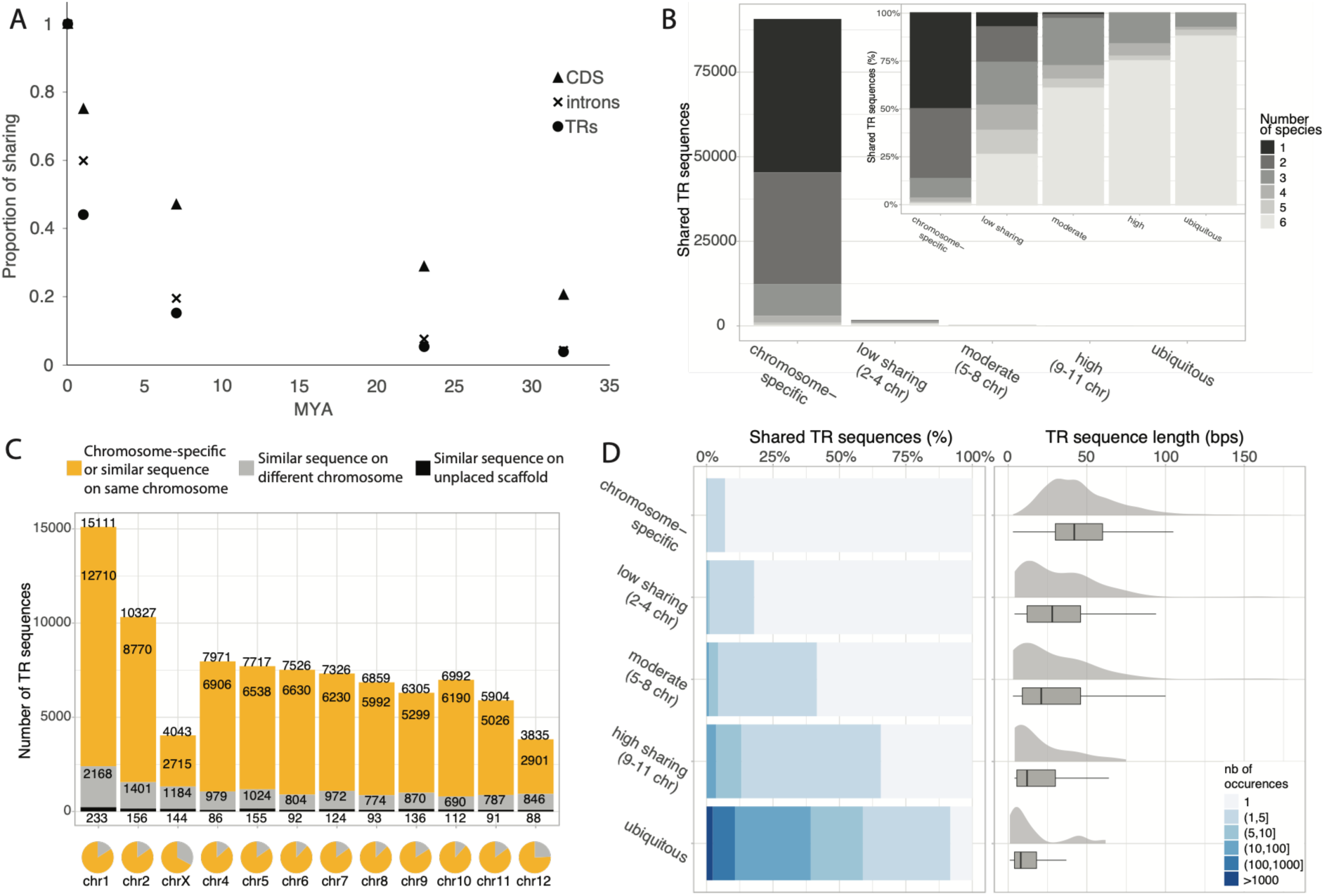
TR sequences in T. poppense display rapid turnover and strong chromosome specificity. (A) Relationship between the proportion of T. poppense sequences shared with species of different divergence times (in million years, MYA), across TRs, coding sequences and introns. (B) Number of unique TR sequences shared between chromosomes (from chromosome-specific to ubiquitously shared across all chromosomes), with grey shading indicating the number of Timema species in which they were identified (inset shows relative proportions). (C) Distribution of TR sequences across the 12 chromosomes of T. poppense, categorized by chromosome specificity. Yellow: TR sequences that are chromosome-specific or share ≥90% sequence similarity with another TR on the same chromosome; grey: TR sequences with a similar sequence on a different chromosome; black: TR sequences with a similar sequence on an unplaced scaffold. Numbers indicate counts per category. Pie charts illustrate the proportion of each category per chromosome. (D) Number of occurrences per chromosome (i.e. number of TR arrays) for each category of TR sequences shared between chromosomes (left panel). Distributions of TR sequence length based on the shared number of chromosomes (right panel).

To further evaluate the selective pressures operating on the TR repeatome, we compared the sequence conservation among *T. poppense* chromosomes, with the assumption that shared sequences may also reflect functional TRs as for chromocentric or telomeric TRs (18, 37, 38). By quantifying the proportion of shared sequences between chromosomes in *T. poppense*, we found that the vast majority of TR sequences (>97%) are chromosome-specific (Figure 3B). Furthermore, these chromosome-specific TRs are largely specific to *T. poppense* (∼50%) or conserved only in the closest sister species *T. douglasi* (∼35%), indicating that limited interspecies sharing overlaps with high chromosome specificity (Figure 3B). We also considered the possibility that our stringent sequence matching criteria (i.e., exact sequence match) may have overestimated the degree of chromosome specificity. To account for this, we calculated pairwise distances among TR sequences in *T. poppense* and identified repeats with at least 90% sequence similarity (see Material and methods). Even with these relaxed criteria, only a small fraction of TRs is characterized by similar sequences across different chromosomes. On average, more than 80% of TR sequences on a given chromosome lack similar sequences on any other chromosome (Figure 3C). Together with our findings on TR repeatome turnover, these results underscore a strong tendency for TR sequences to be both chromosome- and species-specific, supporting the view that most TRs turnover rapidly and lack specific functionality at the sequence level.

To identify individual TR sequences that could serve potential functions within the large pool of non-functional TRs, we focused on the small subset (less than 0.1%) of TRs whose sequences are consistently shared across multiple chromosomes (Figure 3B inset). These conserved TRs may either result from the inherent properties of their sequences to form identical TR arrays independently (a type of homoplasy), or from selective constraints that preserve them for specific functions. The former explanation is particularly plausible for short sequences, as they are more likely to independently form TR arrays composed of the exact same sequence, in comparison to longer sequences. To test this, we quantified for each category of shared TR sequence (ranging from chromosome-specific to shared among all 12 chromosomes) their length distribution and how many distinct arrays they form. Our results indicate that homoplasy is an important driver of TR sequence sharing: ubiquitous TRs are primarily composed of short 2- to 10-bp (median size of 8 bp) sequences, and form 2 to 100 separate arrays (with an average of 10 arrays per chromosome) (Figure 3D). By contrast, chromosome-specific TRs range mostly from 20- to 60-bp in length (median size of 43 bp) and typically form a single array (Figure 3D). Nevertheless, we also find that shared TRs are disproportionately located in TR-poor genomic regions, rather than being randomly distributed along chromosomes, as would be expected under homoplasy being the sole driver of TR sequence sharing (Supplementary figure 3). Taken together, shared TR sequences are likely mostly consequences of homoplasy but a functional role for at least some of them cannot be excluded without additional comparative and/or functional analyses.

Overall, our characterization of TR sequence diversity and genomic distribution within and between *Timema* species reveal a clear pattern in the TR repeatome: a vast majority of TRs are chromosome-specific, exhibit rapid sequence turnover among species and consist of relatively long sequences that occur in single arrays. By contrast, a small fraction of TRs is broadly shared across chromosomes and species. These TRs are generally short sequences that form multiple arrays typically located in TR-poor genomic regions.

### TR abundance and centromere status jointly shape local genomic divergence

Whether TR repeatome turnover occurs uniformly across the genome or varies among genomic regions remains unknown. In particular, centromeres represent a peculiar region to contrast with the rest of the genome as they are typically enriched in tandem repeats, including in *Timema* (39), and are expected to undergo rapid sequence turnover (26). Despite this rapid divergence among species, centromeres perform a highly conserved role in chromosome segregation during cell division, a phenomenon known as the "centromere paradox". The leading explanation for this paradox, the centromere drive hypothesis, proposes that centromeric variants that bias their transmission toward the egg during female meiosis can rapidly spread through populations, thereby accelerating sequence turnover (26). However, rapid centromeric sequence turnover could also simply reflect the generally fast evolutionary dynamics of TRs, as highlighted in our previous results, rather than a centromere-specific evolutionary process. Distinguishing between these alternatives requires comparing the evolutionary dynamics of centromeric TRs with those of TRs elsewhere in the genome.

To identify centromeric regions in *T. poppense*, we performed CenH3-directed chromatin immunoprecipitation (ChIP) and mapped ChIP and input reads to the reference genome (Figure 4A). We then used two complementary approaches to compare the divergence of centromeric and non-centromeric regions in *T. poppense* from its closest sexual relative, *T. californicum* (see details in Material and methods). First, we compared coverage ratios of *T. californicum* and *T. poppense* short reads mapped to the *T. poppense* genome (increased divergence in this approach will be reflected by lower coverage ratios due to reduced mapping quality). Second, we conducted whole-genome alignments between the two species and quantified the fractions of aligned bases across centromeric and non-centromeric regions in *T. poppense*. If centromere drive is at play and accelerates genetic divergence, we expect centromeric regions (particularly TR-rich centromeres) to feature consistently lower values for these two metrics. Because the effect of local TR abundance on divergence is unknown and likely influences the comparison between centromeric and non-centromeric regions, we further included the proportion of TRs within 250-kb non-overlapping genomic windows as a covariate in our analyses.

**Figure 4:**
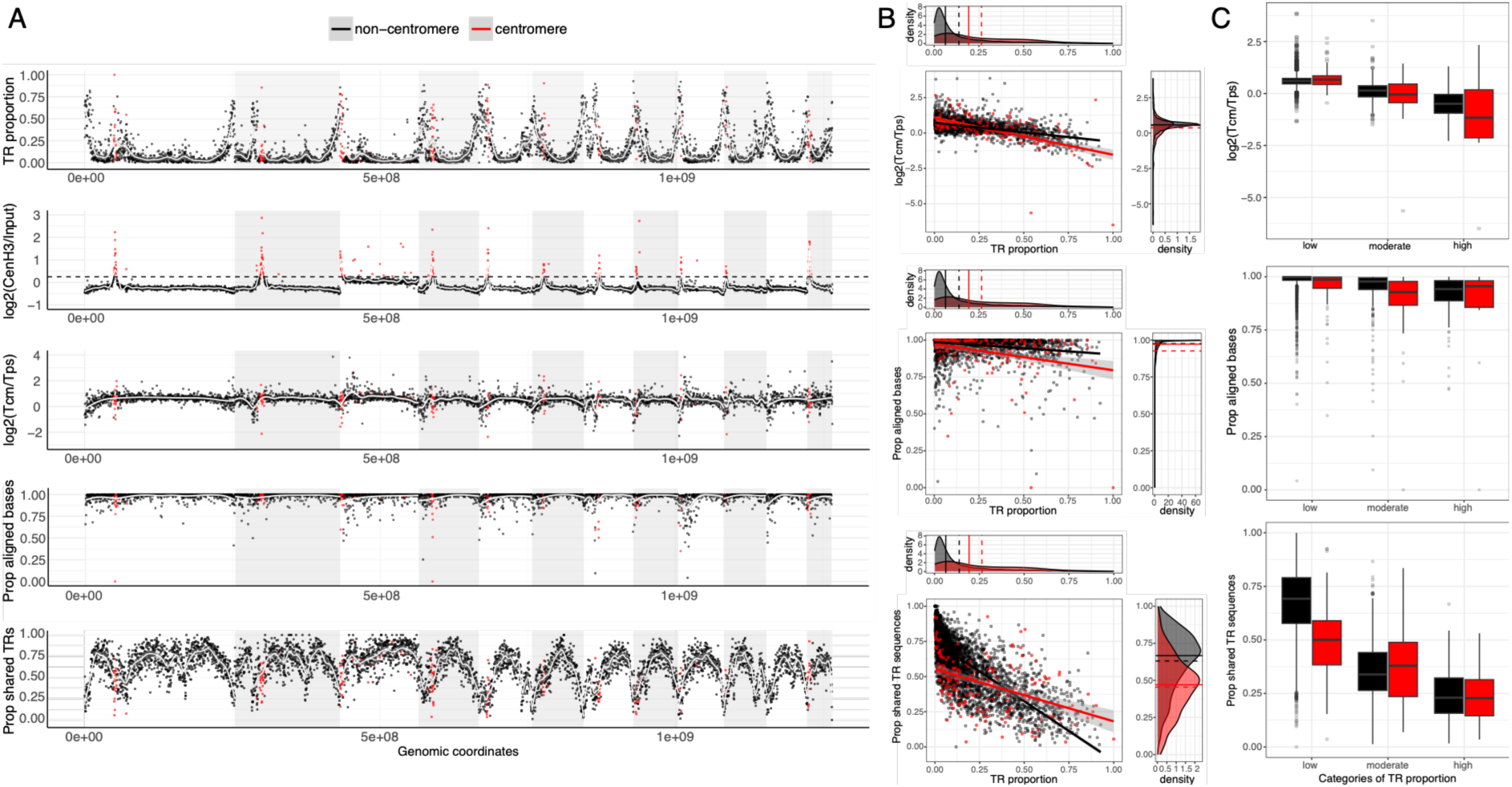
Evolutionary dynamics of centromeric and non-centromeric regions across the twelve chromosomes of T. poppense. Each point represents a 250-kb non-overlapping genomic window, classified as centromeric (red) or non-centromeric (black) based on the CenH3-directed ChIP-seq signal (see Material and methods). Chromosome-wide distributions of, from top to bottom: TR proportion, CenH3 ChIP-seq signal [log₂-transformed of CenH3 versus input control reads], as well as comparisons between T. californicum and T. poppense showing coverage ratios (log2-transformed), proportion of aligned bases, and proportion of shared TR sequences. Alternating white and grey backgrounds indicate individual chromosomes. Centromeric windows are defined by log₂(CenH3/input) > 0.25 (dashed line). For visualization purposes, the lowest coverage ratio values on chromosomes 1 and 4 are not shown. Scaling relationships between local TR proportion and three metrics of divergence: coverage ratios (top), proportion of aligned bases (middle) or proportion of shared TR sequences (bottom). For each relationship, TR proportion distributions for centromeric (red) and non-centromeric (black) regions are shown above the main plot, with the corresponding divergence metric distributions displayed to the right. (C) Boxplots showing three metrics of divergence binned into genomic regions with low, moderate, and high TR proportions, illustrating the effects of TR proportion and centromere status.

For both metrics, we find that divergence increases strongly and significantly with local TR abundance (i.e., TR proportion), independently of centromere status. Both coverage ratios and proportions of aligned bases decrease markedly as the local TR proportion increases in centromeric and non-centromeric regions (Figure 4; Supplementary table 2). This result indicates that the elevated divergence observed in centromeric regions of many species is at least partly attributable to their enrichment in TRs, including in *Timema* (Figure 4). Our analyses also reveal a significant interaction between centromere status and TR abundance for divergence, with centromeric regions becoming more divergent than non-centromeric regions as TR abundance increases (Figures 4B and 4C; Supplementary table 2). This suggests that centromeric TRs evolve more rapidly than non-centromeric TRs in TR-rich genomic regions. Consistent with this interpretation, analyses restricted to TR-annotated sequences alone similarly reveal lower coverage ratios and proportions of aligned bases in centromeric TRs relative to non-centromeric TRs (Supplementary figure 4; Supplementary table 2). These findings are robust with respect to sample size differences: we obtained similar results both genome-wide and when randomly subsampling non-centromeric windows to match the TR proportion distribution observed in centromeric windows (1000 iterations, Supplementary figures 5 and 6; Supplementary table 2).

Overall, our results indicate that the rapid divergence of centromeric regions stems from a combination of two processes: the intrinsically fast-evolving nature of TRs and additional centromere-specific evolutionary forces, possibly related to centromere drive.

### Centromeres are composed of more TR copies within arrays and more similar arrays

Our results above show that TR-rich regions diverge rapidly in both centromeric and non-centromeric genomic contexts. However, whether such rapid evolution is associated with similar or distinct molecular processes is unknown. To uncover putative differences, we first compared the TR organization between centromeric and non-centromeric windows, quantifying array number and size, as well as TR copy numbers and TR sequence length within arrays, while controlling for local TR abundance in *T. poppense*. Centromeric and non-centromeric windows differ markedly in both number and average TR array sizes. As TR abundance increases, centromeric windows harbor fewer but substantially longer TR arrays than non-centromeric windows (Supplementary figure 7; Supplementary table 2). Arrays are longer mostly because they are composed of more TR copies rather than longer sequences (Supplementary figure 7; Supplementary table 2). Thus, centromeric and non-centromeric windows appear to evolve through distinct molecular processes: centromeres are TR rich because of many TR copies within arrays, while non-centromeric windows are TR rich because of multiple TR arrays.

In a second approach, we compared the conservation in TR sequences between centromeric and non-centromeric windows, quantifying across genomic windows the proportion of shared sequences between *T. poppense* and *T. californicum* (hereafter “sequence-sharing metric”), while controlling for chromosome identity and centromere status (see Material and methods). Consistent with our previous analyses, we find that the proportion of shared sequences decreases significantly with increasing local TR abundance in both centromeric and non-centromeric regions (Figure 4; Supplementary table 2). These findings support the general view that rapid local divergence is partly driven by accelerated repeatome turnover in TR-rich genomic regions, including centromeres.

However, centromeric TRs retained generally higher sequence conservation than non-centromeric TRs in regions with high TR abundance. This is evidenced by higher proportions of shared TR sequences in centromeric compared to non-centromeric regions at high TR proportions, whereas the opposite pattern is observed at low TR proportions (Figures 4B and 4C; Supplementary table 2). These conclusions are also supported by our subsampling procedure matching the TR proportion distribution between centromeric and non-centromeric windows (Supplementary figure 6; Supplementary table 2). By comparing the number of TR arrays composed of shared versus *T. poppense*-specific sequences, we show that the elevated sequence conservation observed in centromeres is mainly caused by a marked reduction of arrays with *T. poppense*-specific sequences, particularly in TR-rich regions (Supplementary figure 8A; Supplementary table 2).

This pattern was further confirmed by extending the sequence sharing analysis to the four additional sexual *Timema* species (i.e., across the genus). Specifically, for each annotated TR sequence in *T. poppense*, we quantified the number of *Timema* species comprising the exact same sequence, irrespective of genomic location (values thus ranging from 1 if only present in *T. poppense* to 5 if shared across all species; Supplementary figures 9 and 10; see Material and methods). Averaged across 250-kb genomic windows, these species-sharing values reveal that centromeric TRs display greater sequence conservation (i.e., higher species-sharing values) than non-centromeric TRs in TR-rich regions, owing to their reduced numbers of *T. poppense*-specific arrays (Supplementary figures 8B and 11; Supplementary table 2).

Collectively, these results indicate that TR-rich centromeric and non-centromeric regions follow distinct evolutionary trajectories despite exhibiting similarly elevated levels of divergence. The divergence of non-centromeric TR-rich regions is associated with extensive TR sequence turnover owing to the accumulation of species-specific TR arrays, whereas divergence in centromeric regions appears to involve a larger contribution from copy-number changes or structural reorganization of shared TR sequences. Consistent with this interpretation, non-centromeric regions harbor a greater diversity of TR arrays, measured as the number of arrays with distinct TR sequences per genomic window, than centromeric regions (Supplementary figure 11).

## Material and methods

Genome assemblies and sequenced samples used for TR annotation and proportion estimates We used previously published reference genomes from *T. poppense* (accession number: JBJCJD000000000), *T. douglasi* (JBEUWW000000000), *T. californicum* (JBOIUE000000000), *T. cristinae* (JBOIUC000000000), *T. podura* (JBOIUD000000000) and *T. bartmani* (JBOZRC000000000) from (39-42). In addition, we used 4 to 7 re-sequenced females per species from a previously published dataset (43), available in the NCBI Sequence Read Archive under BioProject accessions PRJNA371785 and PRJNA670663.

### Recombination estimates along *T. poppense* chromosomes

We used LD-bases estimations of the recombination landscape for *T. poppense.* To obtain these estimates, we collected 20 new individuals in the field in 2018 (location 36°99’64.3’’N 121°71’77.8’’W). DNA extractions, library construction and whole genome sequencing for these individuals were performed by the MGX platform (Montpellier). The PCR-free libraries were pooled, and each multiplex was sequenced twice on a separate lane of an Illumina Novaseq6000 S4 flow cell using 150 bp paired-end reads, to ensure a sequencing depth of at least 20x per individual. The raw demultiplexed reads were adapter trimmed and processed for quality control using Fastp (v0.12.4).

We then performed variant calling using the best-practise pipeline in GATK (v4.1.8.1, (44, 45)). First, read mapping was achieved using bwa mem algorithm (v0.7.17, (46)). Mapped reads yielded an average depth coverage per sample of 16.3x and average fraction of successfully mapped reads of 98.8%. We tagged shorter split hits as secondary when multiple primary alignments were generated per query sequences for Picard compatibility (-M option), and -PCR duplicates were marked with Picard MarkDuplicates (-VALIDATION STRINGENCY LENIENT option). Picard AddOrReplaceReadGroups was used to assign reads to a new read-group ID. We then called variants using the HaplotypeCaller, with allele-specific annotations, generating GVCF files (options -G StandardAnnotation, -G AS_StandardAnnotation, -G StandardHCAnnotation). Joint genotyping (GenotypeGVCFs, default settings) was performed on a GenomicsDB workspace, created with GenomicsDBImport, by pooling all individuals for each species. We called 30,741,545 SNPs.

We discarded low-quality SNPs, by applying several filters. We removed SNPs within 5 bp of an indel with Bcftools (v 1.9, (47), -g 5). We used Vcftools (> v 0.1.16, (48)) to exclude indels (--remove-indel), non bi-allelic variants (--max-alleles 2 --min-alleles 2), and SNPs with more than 10% of missing genotypes (--max-missing 0.9). We removed sites with average depth coverage outside the 90th confidence interval of read depth coverage across individuals to control for poorly sequenced regions or duplicated loci (i.e., 4-23x). We applied a Hardy-Weinberg exclusion filter with a p-value threshold of 0.01 (--hwe 0.01). Finally, we removed singletons, which are not informative to infer recombination rates, by applying a MAC (Minor Allele Count) of 1 (--mac 2). After filtering, we retrieved 17,037,010 SNPs.

Haplotypes were phased in two successive steps. First, we used the physical phase information from the individual paired-end reads spanning neighboring heterozygous positions using WhastHap (v1.4, (49)). This resulted in 86.6% of the heterozygous positions being pre-phased. Pre-phased haplotype blocks were then statistically phased at the chromosome level using SHAPEIT4 using default settings (v4.2.2, (50), phase-set error rate: --use-PS 0.0001; and MCMC iteration scheme: 5b,1p,1b,1p,1b,1p,5m). According to the average recombination rate in insects estimated by (51), we assumed a constant recombination rate of 10 cM/Mb, and we used the effective population size estimated from the nucleotide diversity computed in 100-kb windows with Vcftools as input parameters.

We inferred the ancestral allelic states of each variant using the maximum likelihood method implemented in est-sfs (v2.04, (52)). Variants were orientated using the information of three outgroups. We used the previously published reference genome of *T. douglasi* (GCA_901482245.1), *T. californicum* (GCA_902141385.1) and *T. shepardi* (GCA_902151425.1) to orient the SNPs of *T. poppense* (43). We retrieved the 100 bp-flanking sequences of each SNPs in the ingroup species with the getfasta function of Bedtools (53) and used blastn to find their orthologous positions on reference genome of each outgroup species, retaining only the best hits (-outfmt 6, -max_target_seqs 1, -max_hsps 1). Ancestral state probabilities were then inferred with est-sfs from the ingroup allele frequencies and the allelic states of the outgroups. This method also takes into account the phylogenetic relationship between the ingroup and the outgroups that we deduced from (54) and from the mean sequence identity scores computed with blastn. Est-sfs was run using the Kimura-2-parameter substitution model. We estimated population recombination rates *⍴ (⍴*=4*Ner*, where *Ne* is the effective population size and *r* is the recombination rate in M/bp) between successive SNPs using the pairwise composite likelihood method implemented in LDhelmet (v1.19, (55)) for each population independently. We first converted VCF files into fasta sequences using the vcf2fasta function of vcflib (https://github.com/vcflib/vcflib), and into the position and SNPs input format with the --ldhelmet option of VCFtools. We used the probability P and 1-P of the major and minor allele being ancestral computed by est-sfs as prior for the ancestral allelic states. For each chromosome, we used the recommended window size of 50 SNPs to create the haplotype configuration files. The likelihood look-up tables were computed using the recommended grid for the population recombination rate (*ρ*/pb) (i.e. *ρ* from 0 to 10 by increments of 0.1, then from 10 to 100 by increments of 1), and with the Watterson *θ* = 4*Neμ* parameter of the corresponding chromosome computed in 100kb windows with Vcftools and using *μ*=10^-8^. Eleven Padé coefficient tables were computed. The Monte Carlo Markov chain was run for 1 million iterations with a burn-in period of 100,000 and a window size of 50 SNPs, using a block penalty (BP) of 5. We estimated a transition matrix following the method in (55). LDhelmet was run five times independently.

The upper 1% of the population recombination rate (*ρ*/pb) estimates were excluded as outliers. For each interval between successive SNPs, the population recombination rate (*ρ*/pb) was multiplied by the corresponding physical interval length to obtain a length-weighted contribution. For each non-overlapping 250-kb window across the 12 assembled chromosomes, the length-weighted mean population recombination rate (referred to as rho values throughout) was calculated as the sum of the length-weighted *ρ* contributions divided by the total physical length represented by the retained intervals.

### GC estimates along *T. poppense* chromosomes

The proportion of GC content was also calculated within non-overlapping 250-kb windows of the 12 chromosomes, using BEDTools (version 2.30.0, command: bedtools nuc -fi <fasta> - bed <bed>). Specifically, we counted the number of G and C bases within each window and divided these values by the total number of A, T, G, C bases in the same window.

### Tandem repeat annotations

Tandem repeats were annotated separately for each species with Tandem Repeat Finder (TRF) v4.09.1 (56), using the parameters matching weight = 2, mismatch penalty = 7, indel score = 7, match probability = 80, indel probability = 10, minimum alignment score = 50, and motif size up to 2000 bp. TRF provides annotations of all TR arrays that harbour at least 1.8 copies. All analyses were performed on TR arrays of at least 5 copies and of motif sizes varying between 1 to 2000 bp.

### Comparison of TR proportion across species, within and between chromosomes

Illumina short-read data were first trimmed using trimmomatic (v0.39). Trimmed reads were mapped to the species-specific reference genome using minimap2 version 2.28 (options: -ax sr). Reads that mapped equally well to multiple locations in the genome were assigned randomly to one of them. Chimeric reads were removed using SA:Z tags, and PCR duplicates were eliminated with Picard (v2.26.2). We estimated the genome-wide TR proportion for each individual by calculating the ratio of the per-base depth coverage between TR-annotated bases and the total bases in the genome assemblies. The per-base depth coverage was estimated using the *samtools depth* command. TR annotations were used to estimate the depth coverage of the TR-annotated bases. These coverage-based estimates were very similar to TR proportion estimates calculated from the TR array length estimates in the TRF output (see Rscript *Script_TR_proportion_alongChromosomes_coverage-vs-assembly_estimates_TRpaper.R*, available at [GitHub repository](https://github.com/WilliamToubiana/Git_TandemRepeat/tree/main/Rscripts_TR_evolution_paper2)), revealing a strong correlation (correlation coefficient of 0.88, p-value =9.166e-05) between the two methods (Supplementary figure 12).

Similarly, we estimated the proportion of TRs within non-overlapping 250-kb windows by summing the depth coverage separately for TR-annotated bases and total bases within each window and computing the ratio of these sums. We also applied this method at the chromosomal level by aggregating the coverage values of TR-annotated and total bases across all windows of each chromosome.

### Genome size and TR abundance estimates across *Timema* individuals

Genome size for each individual was estimated by dividing the total number of back-mapped bases by the mean per-base coverage. Similarly, TR abundance was calculated by dividing the number of back-mapped TR-annotated bases by the mean per-base coverage of TR-annotated regions. These values were then used to assess how much of the variation in genome size could be explained by TR abundance, by fitting a linear model (genome size ∼ TR proportion) and extracting the R² value.

### Determination of TR repeatome turnover in *T. poppense*

To estimate the turnover of the TR repeatome, we searched for exact matches among TR sequences across *Timema* species. For this, we first generated a non-redundant list of TR sequences that account for redundancies introduced by TRF annotation. TRF assigns a consensus TR sequence to each annotated TR array in the genome assembly; however, some are classified as distinct despite being effectively redundant. These redundancies fall into four main categories: 1) sequences with different starting positions (e.g., *ATGC* and *TGCA*), 2) reverse complement sequences (e.g., *ATGC* and *GCAT*), 3) sequences containing intrinsic tandem repetitions (e.g., *ATGC* and *ATGCATGC*), 4) sequences exhibiting a combination of the above categories. To remove these redundancies, we implemented a custom pipeline that assigns a representative sequence to each annotated TR, based on minimal length and alphabetical order, including reverse complements (Perl script *script_minimal_rotation_parse.pl*, available at GitHub repository). Thus, the above consensus TR sequences (i.e., *ATGC*, *TGCA*, *GCAT* and *ATGCATGC*) would be assigned the representative sequence *ATGC*, because it is the shortest and alphabetically first sequence. We applied this pipeline to all TR annotations before analysing the turnover of the TR repeatome. For the focal species *T. poppense*, we further calculated the total number of arrays, and the average length and copy-number of representative TR sequences per chromosome and per non-overlapping 250-kb windows.

To assess the rate of TR repeatome turnover in *T. poppense*, we conducted pairwise comparisons, counting the number of representative TR sequences in *T. poppense* that had an exact match with those annotated in another *Timema* species.

### Determination of CDS and intron sequence turnovers in *T. poppense*

CDS and intron sequence turnover rates in *Timema* genus were estimated using a whole-genome alignment spanning 30 My of evolution. Eleven assemblies from ten species were aligned using Progressive Cactus (v6.1.0; (57)). Prior to this, assemblies were masked for interspersed repeat using *RepeatModeler* (v2.0.5, with the structural detection of Long Terminal Repeats activated; (58)) and *RepeatMasker* (v4.1.7; (59)), and for TR using Tandem Repeats Finder as described above (v4.09.1). Single-copy CDS and intron sequences were extracted from the alignment using a combination of *cactus-hal2maf* (--dupeMode single – chunkSize 500000) (60) and *maffilter* (v1.3.1; (61)). To estimate turnover rates, sequences were sampled without replacement from the obtained datasets using TR sequence length distribution. For each sequence, it was assessed if it was shared between *T. poppense* and other species, i.e. whether it had a perfect sequence match.

### Determination of TR sequence shared between chromosomes and genomic representation

We searched for representative TR sequences in *T. poppense* that were shared on different chromosomes and represented at different positions along the genome. For this, we counted for each unique representative TR sequence the number of chromosomes on which it occurs and the total number of arrays within and across the 12 chromosomes. Finally, we estimated the proportion of shared TR sequences in 250-kb windows by calculating the ratio between the total number of arrays for the representative TR sequences shared among all 12 chromosomes and the total number of arrays annotated in each window.

### Pair-wise mutational distance between TR sequences annotated in *T. poppense*

We investigated the similarity among representative TR sequences in *T. poppense* by calculating a Levenshtein distance for each pairwise comparison. To facilitate comparisons, all representative TR sequences were size-adjusted by tandem duplications, to a common length defined as the least common multiple of their original lengths, capped at 2000 bp. The resulting distance was normalized by this comparison length to obtain a length-independent dissimilarity score. We then used a custom Python script (*script_minimal_rotation_parse.pl*, available at GitHub repository) to consider all rotations (i.e., all possible starting positions) of one of the two representative TR sequences and retained the rotation that minimized the Levenshtein distance. Pairwise comparisons with at least 90% sequence similarity were kept for downstream analyses to determine the chromosomal locations of the corresponding TR sequences.

### Centromere identification along *T. poppense* and *T. californicum* chromosomes

We determined centromeric regions on *T. poppense* and *T. californicum* chromosomes by identifying genomic regions with enriched CenH3 binding during male meiosis. The histone variant CenH3 has been previously shown to reliably characterize centromeres in *Timema*(39). Specifically, we performed a chromatin preparation on 40 mg of dissected testes immediately frozen in liquid nitrogen in *T. poppense*. Owing to limited sample availability in *T. californicum*, 35 mg of whole male and female bodies were included in the chromatin preparation. CenH3-directed chromatin immunoprecipitation followed by sequencing (ChIP-seq) was then performed as described in (39). ChIP and corresponding input reads were quality-trimmed using trimmomatic (version 0.39) and mapped to the corresponding reference genome using the BWA-MEM algorithm (v0.7.17), with a relaxed seed threshold (- c 1000000000) to maximize alignment sensitivity within highly repetitive regions such as centromeres. Chimeric reads were removed using SA:Z tags, and PCR duplicates were eliminated with Picard (version 2.26.2). Mean coverage was computed for ChIP and input reads within nonoverlapping 250-kb windows across all scaffolds using BEDTools (version 2.30.0) and normalized by the number of mapped reads in each library. Centromeric regions in *T. poppense* were defined as 250-kb windows with a mean ChIP coverage at least ∼20% higher than the mean input coverage [i.e., log2(ChIP/input) ≥ 0.25]. In *T. californicum*, the overall ChIP signal was weaker, resulting in a lower proportion of windows exceeding this threshold. To account for this difference and obtain comparable numbers of centromeric windows, we also applied a more permissive threshold (log₂[ChIP/input] ≥ 0; Supplementary figure 13). Both thresholding approaches led to qualitatively similar results; therefore, only results obtained using the latter criterion are reported here.

### Comparison of coverage ratios from *T. californicum* and *T. poppense* short reads mapped onto the *T. poppense* genome

Trimmed Illumina reads from *T. poppense* and *T. californicum* individuals used for the reference genome assemblies were mapped to the *T. poppense* genome using minimap2 (v2.28, options: -ax sr). Reads that mapped equally well to multiple locations in the genome were assigned randomly to one of them. Chimeric reads were removed using SA:Z tags, and PCR duplicates were eliminated with Picard (version 2.26.2). We then estimated genome-wide and TR coverage for *T. poppense* and *T. californicum* mapped reads, separately, by first calculating the per-base depth coverage of each base and TR-annotated bases, respectively. Specifically, the per-base depth coverage was estimated using the *samtools depth* command and the TR annotation from *T. poppense*. The per-base depth coverages were then averaged within non-overlapping 250-kb windows of the *T. poppense* genome and normalized by the number of mapped reads in each library. Finally, we computed the normalized ratios and log2-transformed the values to compare the coverage ratios between centromeric and non-centromeric genome regions.

### Comparison of the fractions of aligned bases in whole-genome alignments of *T. poppense* and *T. californicum*

The genomic coordinates of all annotated TRs in the *T. poppense* genome were extracted from the repeat annotation. These TR intervals were mapped to the *T. californicum* reference genome using halLiftover from Cactus v2.8.4 (60) based on the whole-genome alignment, keeping only non-duplicated regions (--noDupes parameter). For each non-overlapping 250-kb genomic window, we calculated the proportion of TR sequence that successfully aligned to the orthologous region in *T. californicum*, defined as the total length of aligned TR bases divided by the total TR content in that window. These window-specific aligned proportions were computed using custom scripts (*compute_aligned_proportions.py* and *new_prop_TR_perwindow.sh*, available at GitHub repository).

### Proportion of TR sequence shared between *T. poppense and T. californicum*

Each representative TR sequence in the *T. poppense* genome was assigned a binary sequence-sharing score, indicating whether the representative sequence was shared (1) or absent (0) in *T. californicum*. To be classified as shared, the corresponding TR sequence in *T. californicum* was required to occur on the same chromosome and centromere status (i.e., within a centromeric or non-centromeric region). We then quantified the proportion of shared TR sequences genome-wide, defined as the number of shared representative TR sequences divided by the total number of representative TR sequences for each non-overlapping 250-kb window.

### Genomic landscape of TR sequence conservation among *Timema* species

Each annotated representative TR sequence in the *T. poppense* genome was assigned a species-sharing value based on the number of *Timema* species in which it was identified. Thus, these values range from 1 (specific to *T. poppense* only) to 5 (shared across all five sexual *Timema* species). These values were then averaged within non-overlapping 250-kb windows. Consequently, windows with higher species-sharing values represent genomic regions composed of TRs with greater sequence conservation among *Timema* species.

## Discussion

Ever since the discovery of tandem repeats (TRs) as major components of genomes, their function (or lack thereof) has been debated. Although both functional and non-functional TRs are likely represented in all eukaryote genomes, the lack of genome-wide investigations of TR evolution limits our understanding of TR prevalence and contribution to genome diversification. Our study provides the first in-depth characterization of TR evolution across species and genomic regions while considering the effects of recombination, genetic drift, and selection in the stick insect group *Timema*.

We show that TRs tend to accumulate in *Timema* species with lower effective population sizes, consistent with a non-functional framework that views TRs as slightly deleterious elements and accumulating in genomes when selection becomes less effective (7, 8). Under this framework, TRs are also expected to preferentially accumulate in low-recombination regions owing to locally reduced efficacy of selection. However, we observed the opposite pattern in *Timema*, by which TRs are more abundant in high-recombination regions. Together, we propose that the mutational effects of recombination (or recombination-associated features) on TRs can result in locally elevated TR loads under mutation-selection balance. Two additional lines of evidence support this interpretation. First, high recombination regions are characterized by an increased array density, longer TR sequences, and more TR copies. Second, these regions present an excess of species-specific TR arrays, which, together, suggest that recombination is an important source of *de novo* TR formation in *Timema*.

Simulation models developed and extended in the 90ies, predict that recombination alone can be sufficient (i.e., without the requirement of additional processes as amplification or selection) to account for the presence and accumulation of TRs in the human genome (62, 63). Specifically, these models show that with high rates of recombination, there is a reasonable likelihood of generating a rapid phase of TR accumulation (i.e., TR arrays of at least 20 copies) from an initial TR array of 2 copies, through a random walk of unbiased array expansion and contraction driven by unequal crossover exchange. Under this scenario, high-recombination regions should be favorable for the accumulation of large, potentially hypervariable, TR arrays that would then persist over variable evolutionary timescales.

An alternative to the positive effect of recombination on TR formation would be that the effect of recombination is inherently biased toward TR expansion, as proposed in *Daphnia pulex* (19), or that TRs themselves promote recombination, for example through mechanisms such as increasing double-strand breaks (64). This second scenario may, in turn, generate a positive feedback loop by which TRs would generate increased recombination in TR-rich regions, and consequently favor the formation and expansion of TR arrays by unequal crossing over.

Finally, we cannot rule out the possibility that interspecies variation in recombination rate may also explain variation in TR abundance genome-wide, in a similar manner as it explains the TR variation within genomes. However, intronic GC content (used as proxy for genome-wide recombination rate), is similar across the 5 studied sexual *Timema* species, suggesting that recombination rate remained relatively stable in this group. Taken together, our findings suggest that variation in TR load within and between *Timema*genomes is primarily shaped by genetic drift and meiotic recombination.

The recombination-associated mutational process is also likely to be a major determinant of the TR load variation observed between chromosomes, including the TR reduction on the X chromosome, as the X is expected to recombine less than the autosomes in XX/X0 sex determination systems. This interpretation is further supported by the observed negative relationship between TR proportion and chromosome size, whereby smaller chromosomes, characterized by higher per base recombination rates, harbor higher TR proportions. Extending these between-chromosome comparisons of TR load to other taxa will be crucial to assess whether this relationship represents a general feature of genome evolution.

Conversely, if X-specific reductions in TR content are not detected in other organisms, general sex chromosome-associated processes such as reduced meiotic recombination would be unlikely drivers of the TR reduction on *Timema* X chromosomes. Instead, this would point to a *Timema*-specific pattern. Recent studies of meiotic chromosome behavior in *Timema* have uncovered an unusual pattern in which the centromere-specific histone CenH3 is deposited and maintained along the entire length of the X chromosome throughout male meiosis I, instead of being localized to the functional centromeric region as in autosomes (39). Although the functional significance of this atypical CenH3 distribution on the X remains unclear, it may interact with TR evolution on this chromosome.

Beyond their variation in abundance, TRs are notoriously famous for their rapid sequence evolution; a feature often cited in support of their presumed lack of function. Our comparative analyses of sequence turnover across *Timema* genomes are consistent with this hypothesis and further reinforce the view that the bulk of TRs are non-functional. TRs turnover faster than functional genomic elements such as CDS but also faster than elements typically considered less constrained, such as introns. In addition, the strong chromosome-specificity of TR sequences indicates that they evolve largely independently within genomes, as expected under a predominantly non-functional scenario.

Nevertheless, certain TRs such as centromeric TRs challenge this non-functional view, by exhibiting rapid sequence turnover in spite of their conserved implication in chromosome segregation (26). Our results align with this general observation, revealing that *Timema* centromeres are one of the most rapidly changing regions of the genome, largely composed of TRs with high sequence turnover between species. We further show that this rapid evolution is partly attributable to the fast-evolving nature of TRs *per se* and is consequently a broad property of TR-rich genomic regions. Our analyses also revealed that centromeric and non-centromeric TRs follow distinct evolutionary trajectories, particularly in TR-rich genomic regions: centromeric TRs exhibit higher degrees of divergence than non-centromeric TRs, despite being composed of more TR sequences shared among species. We propose that this pattern results from different architectures of TRs, by which centromeric regions evolve primarily through copy-number changes within arrays and positional rearrangements of shared TRs. By contrast, non-centromeric TR regions evolve more through the formation of species-specific and distinct TR arrays. The processes responsible for these different evolutionary trajectories remain unknown, although differences in the mutational process such as recombination or selective pressure such as centromere drive are key hypotheses to assess in the future.

Overall, our findings show how the rapid divergence and turnover of centromere sequences reflect not only the action of centromere-specific features, but also more general processes related to the intrinsically fast-evolving nature of TRs.

## Acknowledgments

We thank the Lausanne GTF platform for help with ChIP-seq; the Bioinformatics Competence Center for help and discussions on TR sequence turnover; Nicolas Galtier and Pierre-Alexandre Gagnaire for help with the recombination rate analysis; and current and previous members of the Schwander lab for discussions.

## Author contributions

W.T. and T.S. designed the study. W.T. performed molecular work. W.T., M.L., V.M., and J.S. developed methods. W.T. and T.S. analyzed the data, W.T. and T.S. wrote the paper, with input from all authors.

## Data and materials availability

All data needed to evaluate the conclusions in the paper are present in the paper and/or the Supplementary Materials. Raw sequence reads used for genome assemblies have been deposited in NCBI’s sequence read archive BioProject PRJNA1268975. Data were processed to generate plots and statistics using R version 3.4.4. The codes used to run the analyses are available on GitHub (https://github.com/WilliamToubiana/Git_TandemRepeat and https://github.com/WilliamToubiana/Git_TandemRepeat-Rscripts_TR_evolution_paper).

