## Supplementary materials for "Drivers of tandem repeats variation within and between stick insect genomes"

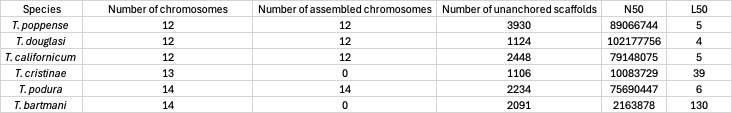

Supplementary table 1: Summary statistics of the genome assemblies (previously published, see Material and methods) used in this study to annotate TRs. The species T. douglasi reproduces via parthenogenesis, the remaining species are sexual. The number of chromosomes refers to karyotype information from (28).

Supplementary table 2: Models and statistics used to compare the evolution of centromeric and non-centromeric TRs in T. poppense.

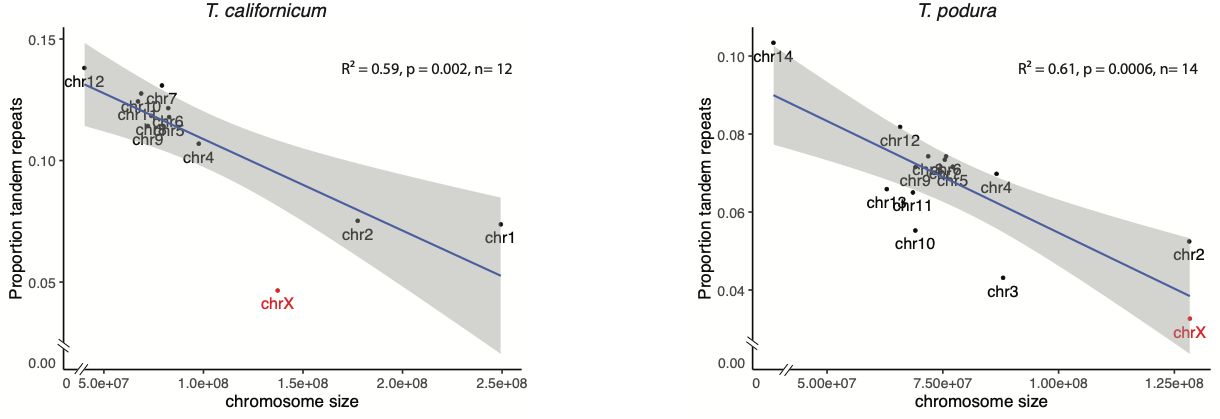

Supplementary figure 1: Relationships between chromosome size and TR proportion in two additional Timema species. Note that T. cristinae and T. bartmani are not included as their genome assemblies are not at the chromosome level. R-squared and p-values were calculated with the X chromosome included.

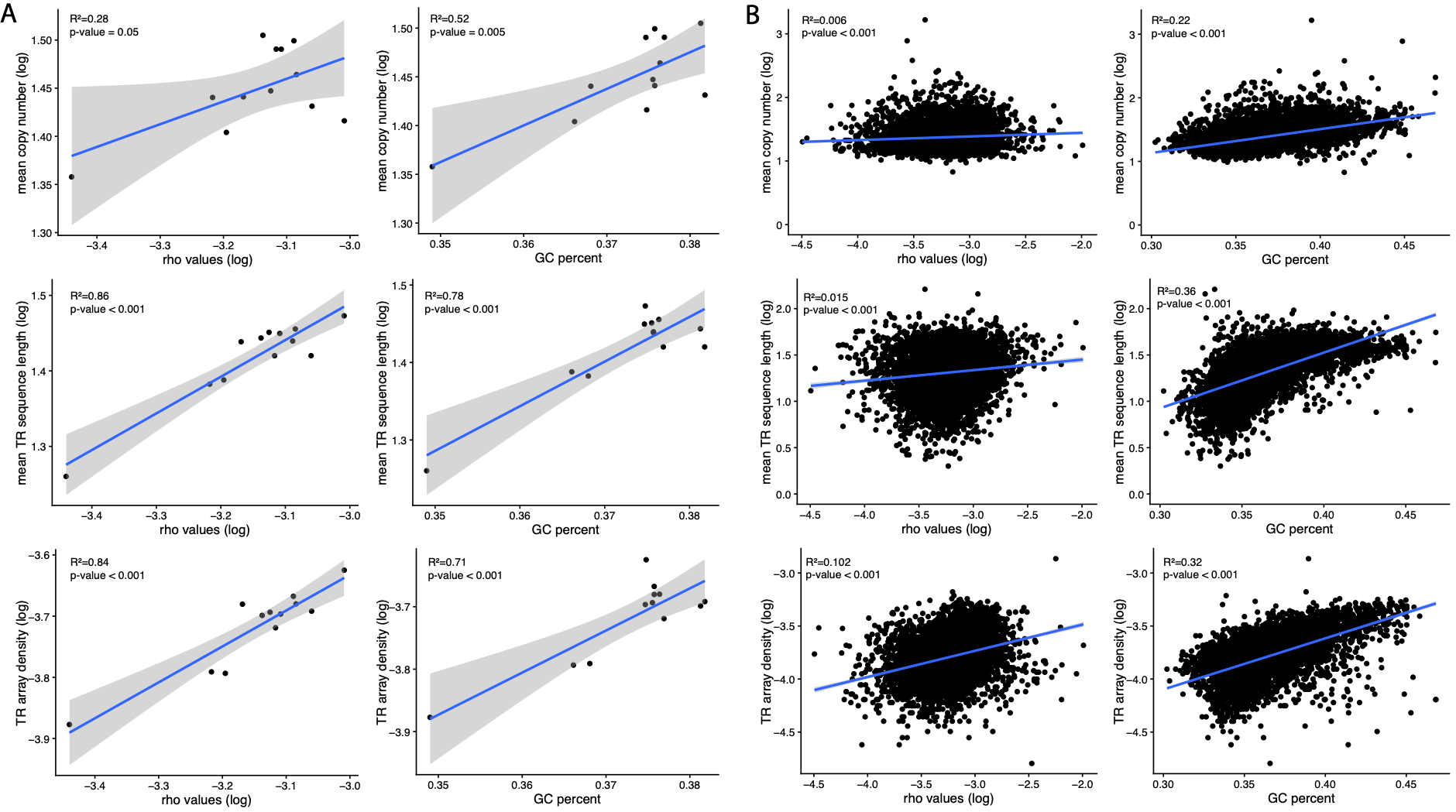
Supplementary figure 2: Relationships between recombination proxies (rho values and GC content) and TR copy number, sequence length and array density across T. poppense chromosomes (A) and 250-kb regions (B).

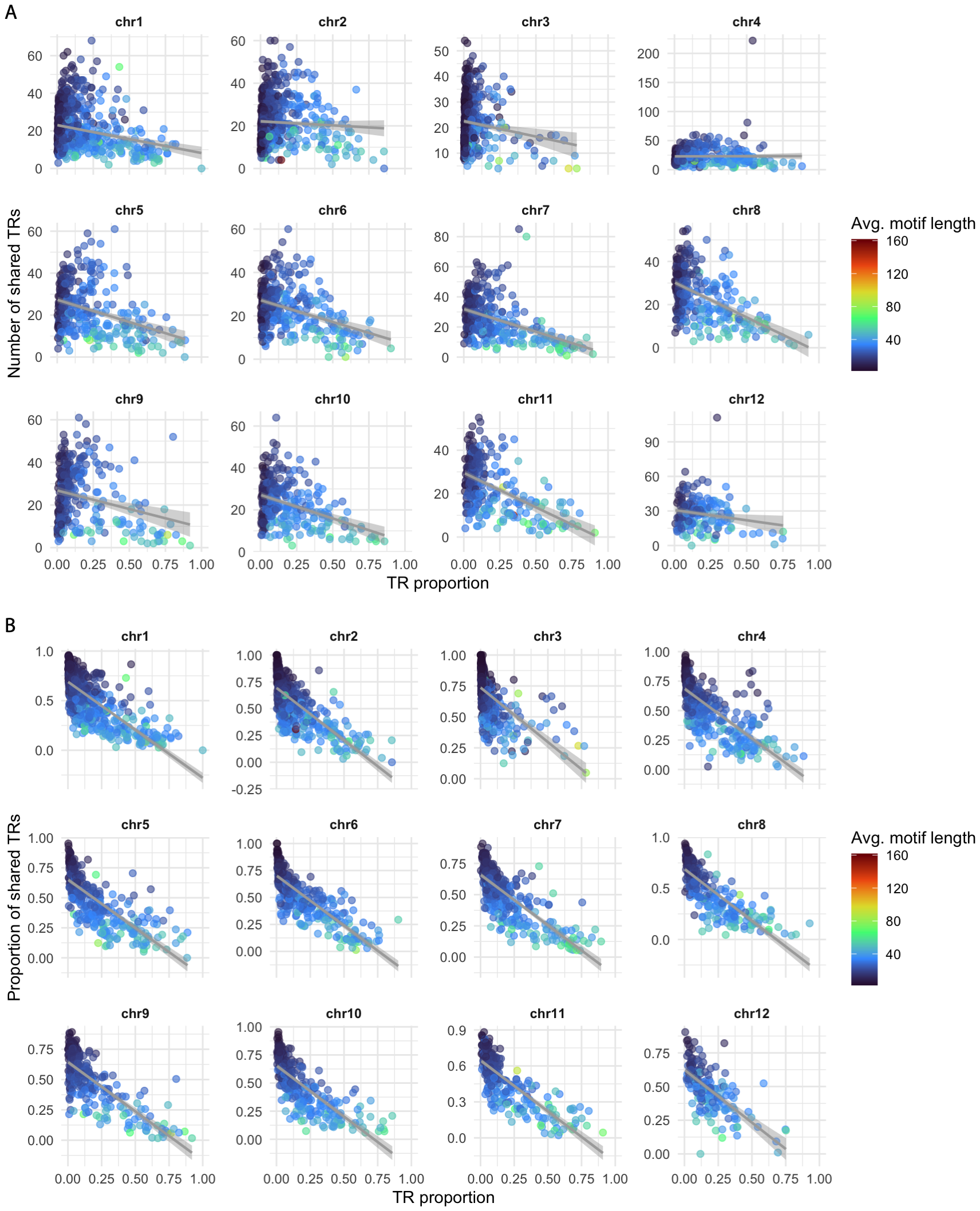
Supplementary figure 3: Number (A) and proportion (B) of shared TR sequences (i.e., sequences shared across all 12 chromosomes of T. poppense; corresponding to the “ubiquitous” category in Figure 3B) for every 250-kb region of a chromosome, with regions color-coded by the mean TR sequence length. The proportion was calculated by dividing the number of shared TR sequences by the total number of TR sequences in the corresponding 250-kb genomic region.

Supplementary figure 4: Comparisons of centromeric and non-centromeric genomic windows when restricting analyses to TR-annotated bases in the genome (A) Chromosome-wide distributions of TR proportion and CenH3 ChIP-seq signal [log₂-transformed of CenH3 versus input control reads]. Note that this panel is the same as Figure 4A, in the main text, but was indicated again here for comparison with panel B. (B) Comparisons between T. californicum and T. poppense showing TR coverage ratios (log2-transformed) and proportion of aligned TRs. Alternating white and grey backgrounds indicate individual chromosomes. Centromeric windows are defined by log₂(CenH3/input) > 0.25 (dashed line) and colored in red, while non-centromeric windows are colored in black. (C) Scaling relationships between local TR proportion and two metrics of divergence: TR coverage ratios (top), proportion of aligned TRs (bottom). For each relationship, TR proportion distributions for centromeric (red) and non-centromeric (black) regions are shown above the main plot, with the corresponding divergence metric distributions displayed to the right. (D) Boxplots showing the two metrics of divergence binned into genomic regions with low, moderate, and high TR proportions, illustrating the effects of TR proportion and centromere status.

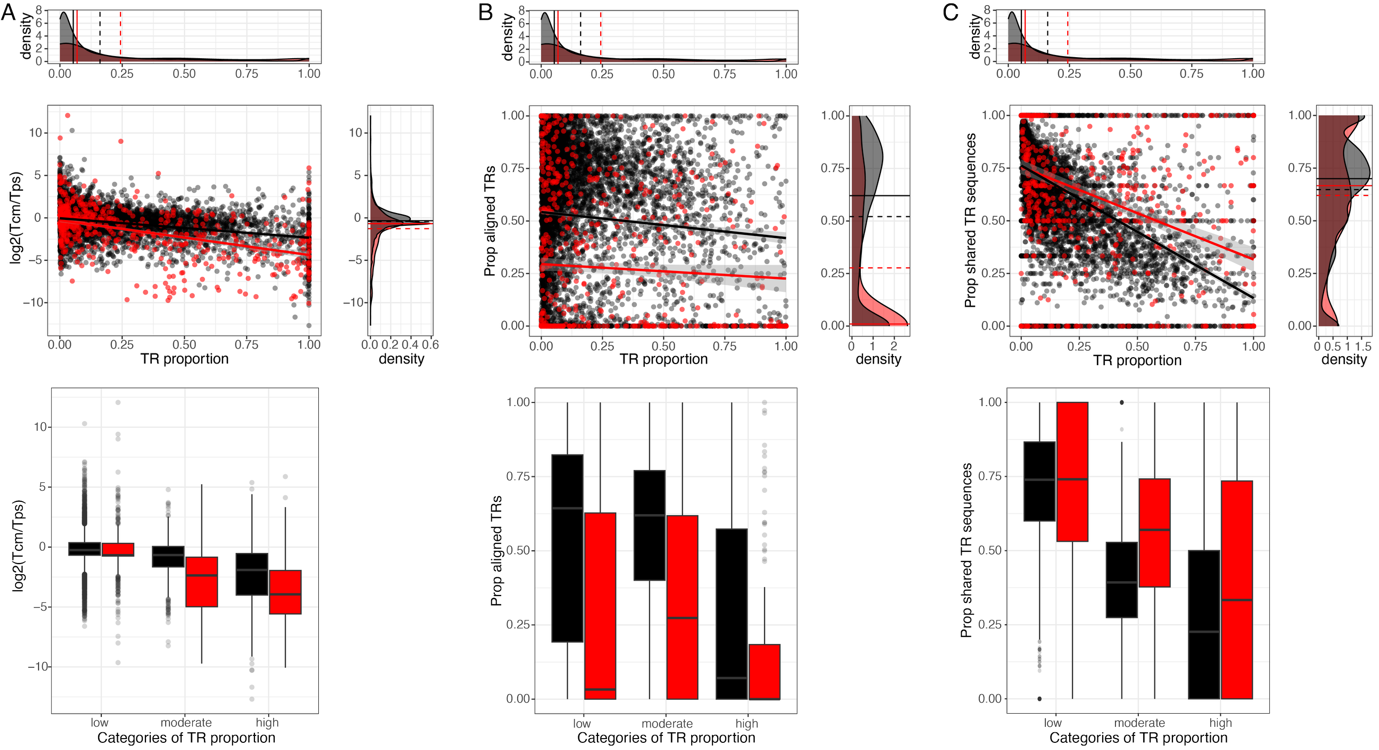

Supplementary figure 5: Genome-wide scaling relationships and boxplots between TR proportion and two metrics of divergence: TR coverage ratios (A) and proportion of aligned TRs (B). Centromeric windows are defined by log₂(CenH3/input) > 0.25 and colored in red, while non-centromeric windows are colored in black.

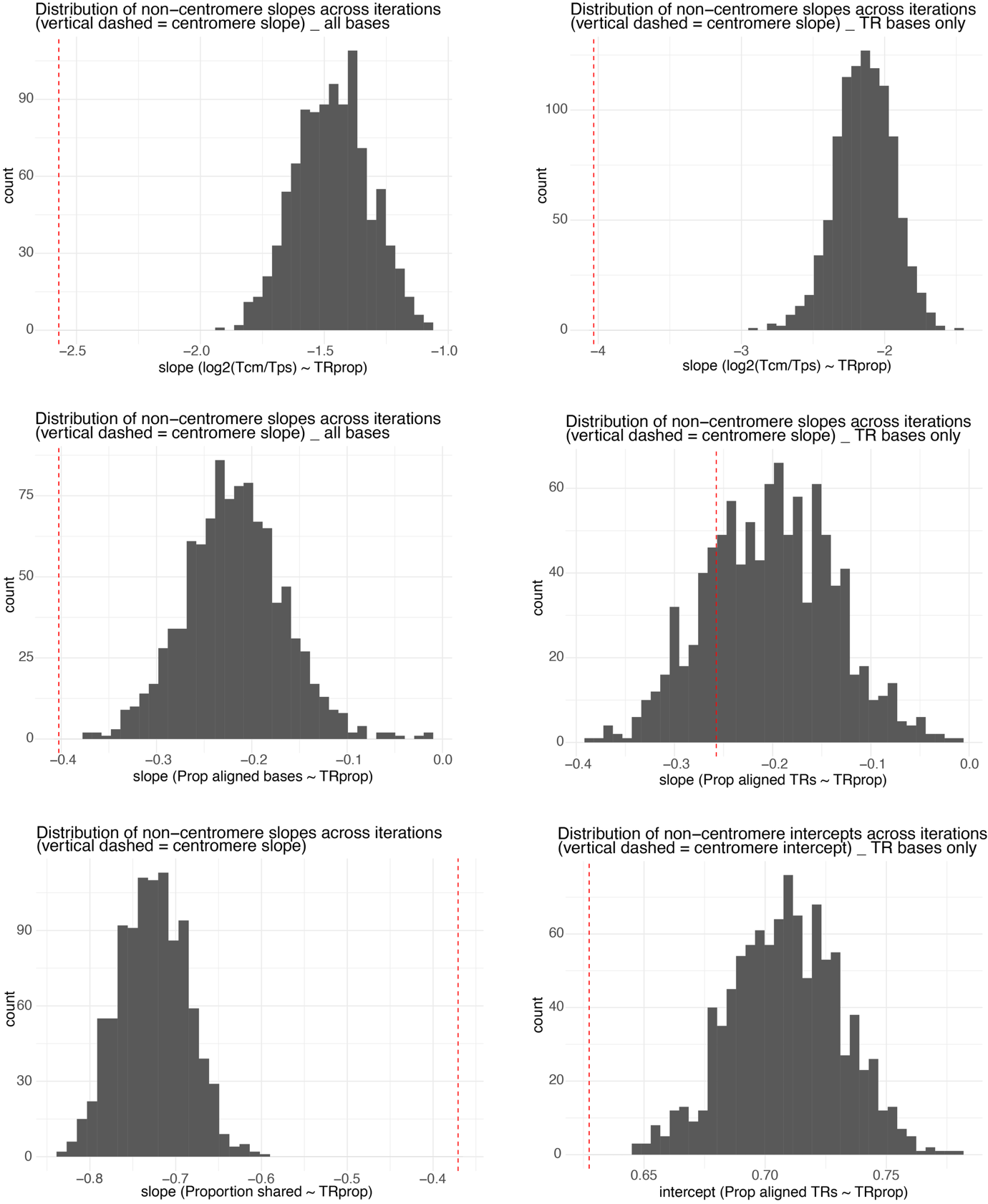

Supplementary figure 6: Distributions of non-centromeric slope or intercept values (for the different metrics of divergence) across the 1000 iterations are displayed. The corresponding centromeric slope or intercept value is indicated by a red dashed vertical line. Note that the terms “slopes” refer here to the negative relationships estimated between TR proportion and the two metrics of divergence (i.e., coverage ratios and proportions of aligned bases), whereas the term “intercepts” refers to the difference in elevation between these relationship estimates.

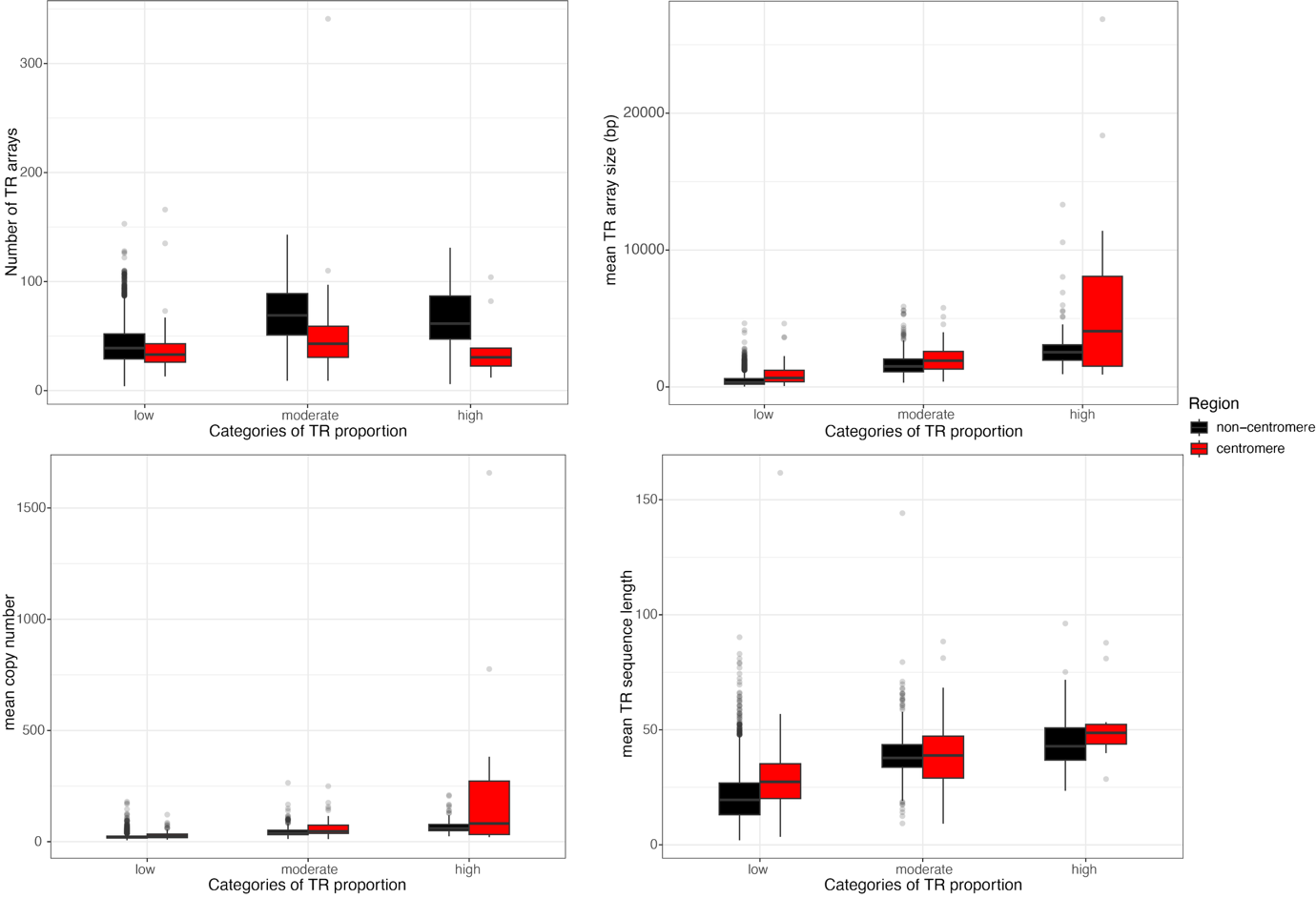

*Supplementary figure 7: Comparisons of centromeric and non-centromeric windows (i.e., 250-kb non-overlapping windows) for the number of TR arrays and the average size of TR arrays (upper plots). The comparison of average TR array size was also decomposed into average copy number and TR sequence length within these arrays (bottom plots).*

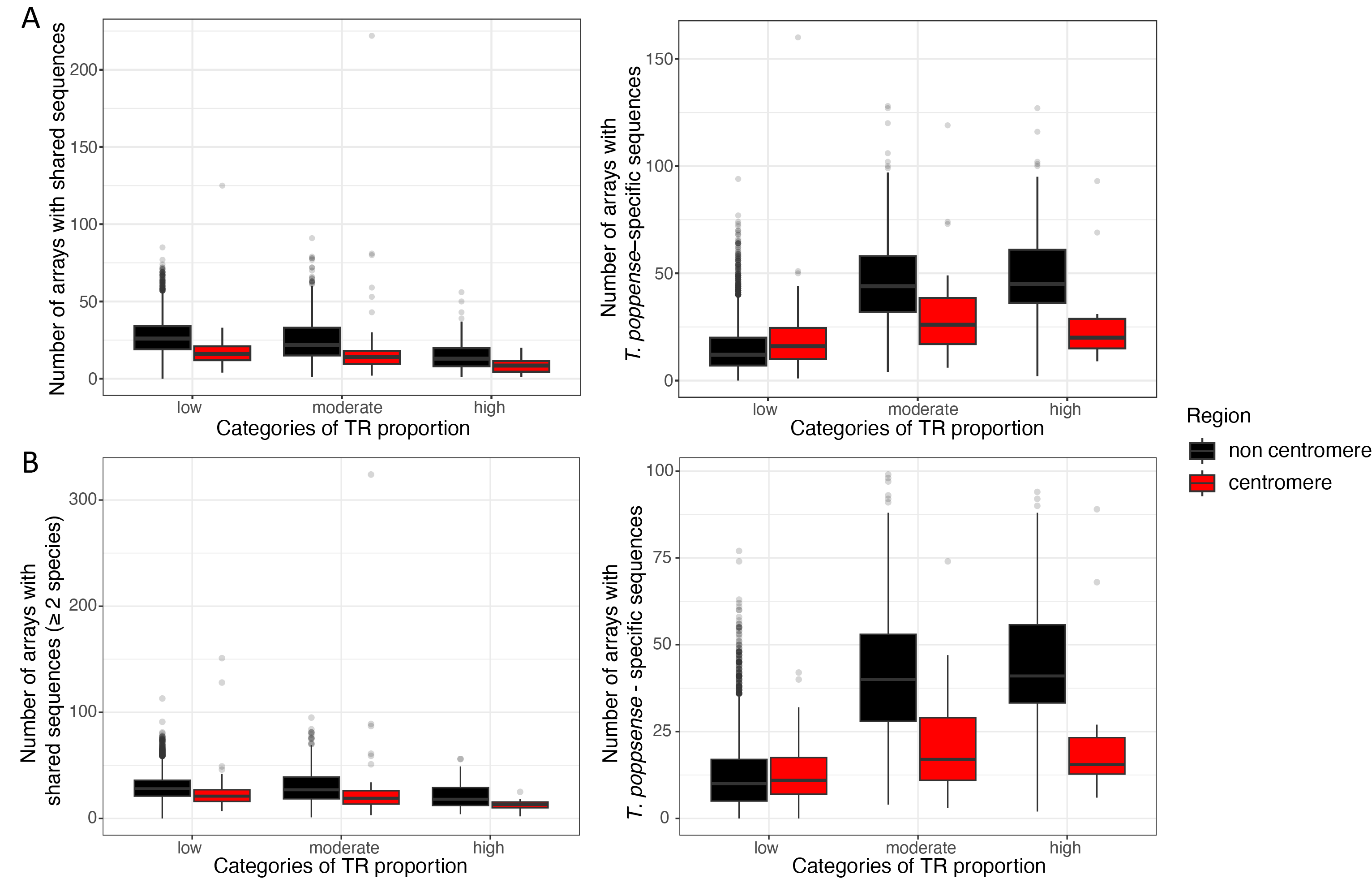

Supplementary figure 8: Number of arrays in T. poppense with shared TR sequences between species and T. poppense-specific TR sequences. The number of shared sequences was quantified either between T. poppense and T. californicum only (A) or across all analyzed sexual Timema species (B). In the latter quantification, we consider a TR sequence as shared when it is identified in at least another Timema species (i.e., ≥ 2 species).

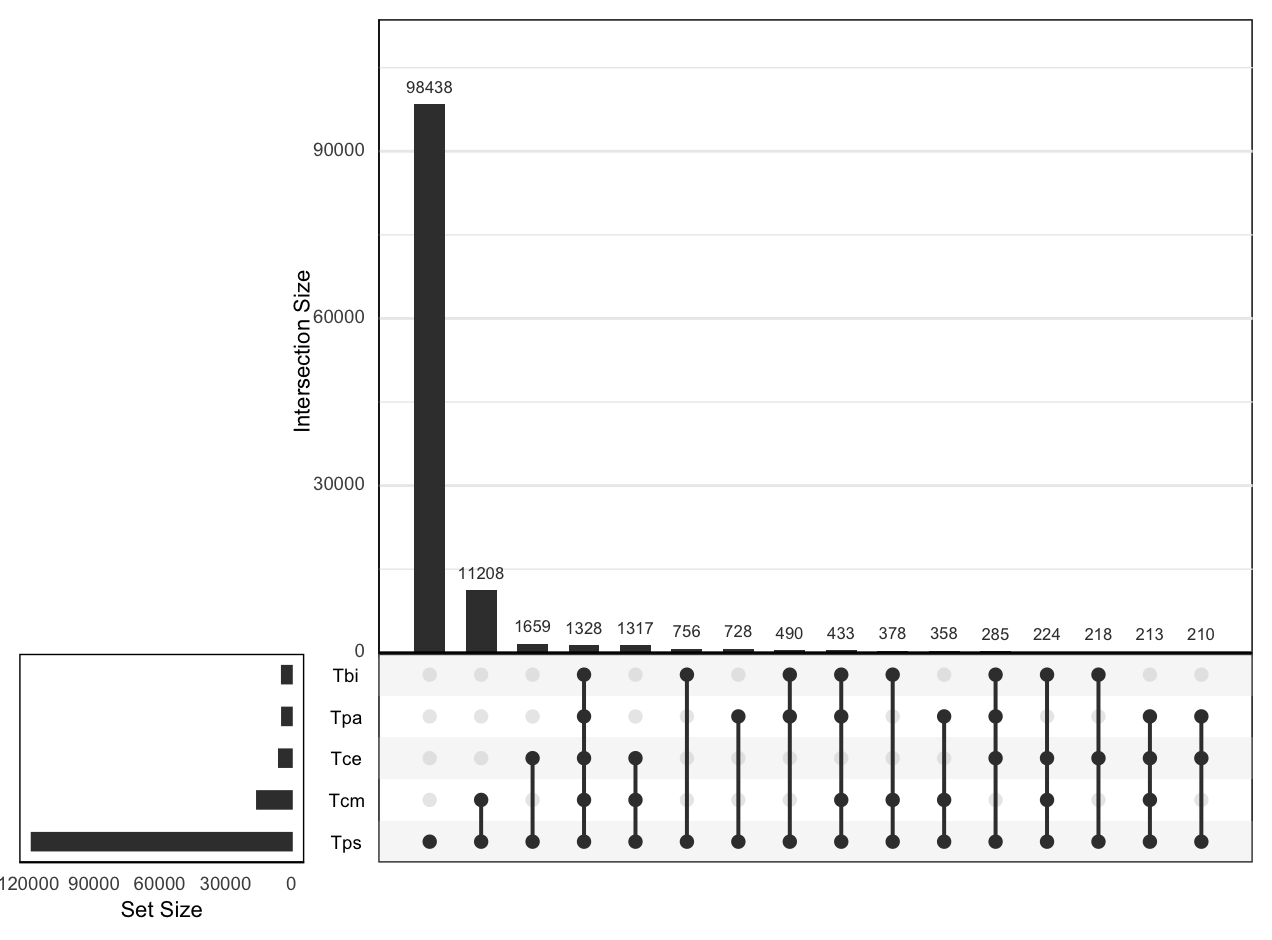

Supplementary figure 9: Intersection of shared TR sequences between T. poppense (Tps) and the four other sexual species: T. bartmani (Tbi), T. podura (Tpa), T. cristinae (Tce) and T. californicum (Tcm).

Supplementary figure 10: (A) Chromosome-wide distributions of, from top to bottom: TR proportion, CenH3 ChIP-seq signal [log₂-transformed of CenH3 versus input control reads] and averaged values of the number of TR sequences shared across the five sexual Timema species (i.e., species-sharing values). Alternating white and grey backgrounds indicate individual chromosomes. Centromeric windows are defined by log₂(CenH3/input) > 0.25 and colored in red, while non-centromeric windows are colored in black.
(B) Scaling relationships between local TR proportion and species-sharing values. For each relationship, TR proportion distributions for centromeric (red) and non-centromeric (black) regions are shown above the main plot, with the corresponding species-sharing value distributions displayed to the right. For each relationship, the distributions of TR proportions in centromeric (red) and non-centromeric (black) regions are shown above the plot, while the distributions of species-sharing values are shown to the right.
(C) Boxplots showing the species-sharing values across genomic regions with low, moderate, and high TR proportions, illustrating the effects of TR proportion and centromere status.

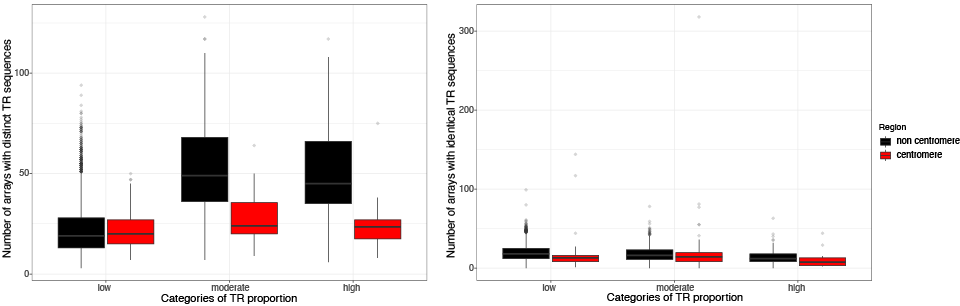

Supplementary figure 11: Number of TR arrays with distinct (left) or identical (right) TR sequences in each 250-kb genomic window.

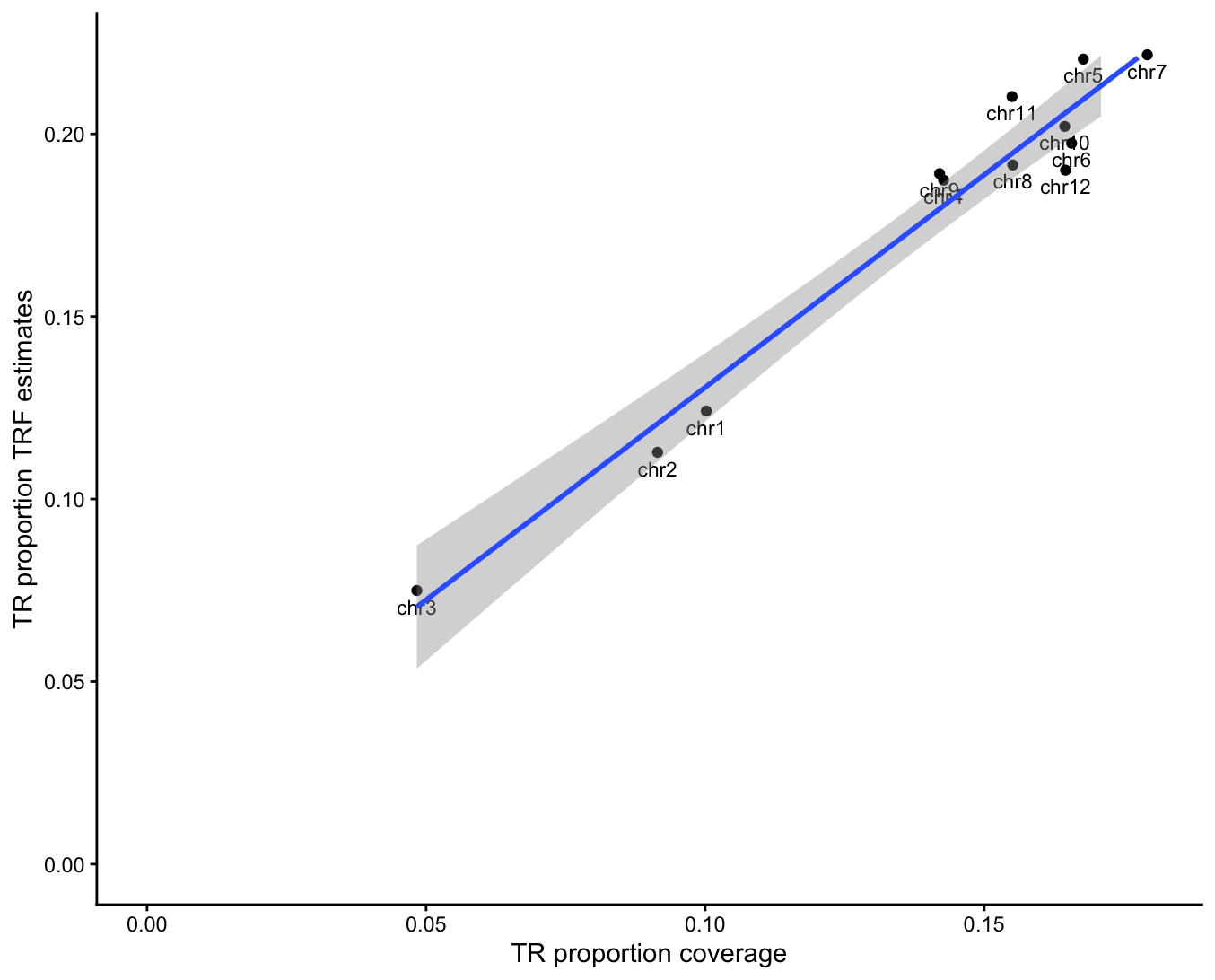

Supplementary figure 12: Relationship between estimated TR proportion from the per-base coverage method and TRF estimates for T. poppense (adjusted R^2^= 0.95; correlation coefficient of 0.89).

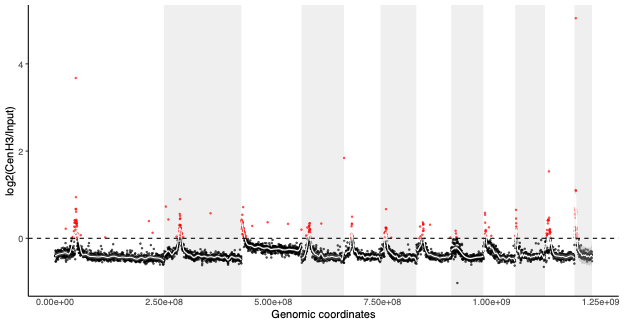

Supplementary figure 13: CenH3-directed ChIP-seq signal in T. californicum [log₂-transformed coverage ratio of CenH3 immunoprecipitation (ChIP) reads versus input control]. The dashed line indicates the threshold (log₂[ChIP/input] ≥ 0) used to define centromeric (red) and non-centromeric (black) windows.
